# Sensitivity to temozolomide of cell lines derived from brain tumours and melanomas: relationship to MGMT gene expression

**DOI:** 10.1101/2024.11.30.626133

**Authors:** Bruce C Baguley, Wayne R Joseph, Pamela M Murray, Kathryn M Stowell, Euphemia Y Leung, Elaine S Marshall

## Abstract

**Background:** We have developed a series of low-passage melanoma and glioma cell lines and are using them to study responses to antitumour drugs. Here we have assessed responses to temozolomide, a commonly used clinical drug used for the treatment of glioblastoma and melanoma. Temozolomide acts mainly by methylation of DNA guanine and lesions can be removed by the repair enzyme O^6^-methylguanine-DNA methyltransferase (MGMT). Low expression of MGMT is one of the main features associated with sensitivity to temozolomide and is usually associated with CpG methylation of the MGMT promoter. We wished to determine whether temozolomide sensitivity in these lines was related to MGMT promoter methylation and MGMT mRNA synthesis. Methods: 63 melanoma and 37 glioblastoma lines were derived from developed from surgical samples, using atmosphere of 5% oxygen for development and propagation in order to minimise oxygen toxicity. Temozolomide sensitivity (IC_50_ data) was analysed by its effect on ^3^H-thymidine incorporation in 5-day assays. MGMT promoter methylation status was assessed by the methylation-specific polymerase chain reaction (PCR). MGMT mRNA expression was assessed using reverse transcription and PCR. Results: 40% of the melanoma lines and 24% of the glioblastoma showed some sensitivity to temozolomide. Of the 79 cell lines analysed for MGMT mRNA expression, a good correlation was found to temozolomide resistance (p < 0.0001). Of the 100 cell lines assessed for MGMT promoter status, 12 showed methylation of both promoter sequences and were all temozolomide-sensitive; 50 lines showed no MGMT promoter methylation and 90% of these were temozolomide resistant; 30 lines showed partial promoter methylation and 53% of these were temozolomide resistant. Conclusion: Temozolomide sensitivity matched MGMT mRNA expression well but only partially matched MGMT promoter methylation. Other factors, such as loss of DNA mismatch repair capacity, may contribute to resistance.

## Introduction

We have developed a series of low-passage tumour lines under an atmosphere of 5% oxygen for development and propagation in order to minimise oxygen toxicity (1). The cell line collection has been analysed extensively in terms of DNA sequence, gene expression (2) and response to sensitivity to anticancer drugs (3). In this study we have assessed the response of these lines to temozolomide, one of the few drugs that are effective in treating glioblastoma (4) that is also in use for the treatment of melanoma. Here we have utilised the collection to gain an increased understanding of its action on cell lines derived from glioblastoma and metastatic melanoma.

Under physiological conditions, temozolomide undergoes spontaneous hydrolysis to MTIC (5- (3-methyltriazene-1-yl)imidazole-4-carboxamide), which methylates DNA predominately at the O^6^-position of guanine (5). While the presence of O^6^-methylguanine in DNA has low intrinsic cytotoxicity, it acts as a recognition point for the assembly of mismatch repair (MMR) complexes that induce stalling of DNA replication forks during the S-phase of the cell cycle (6). These in turn activate kinases such as ATR (Ataxia telangiectasia related complex) (7), leading to G_2_-phase cycle arrest, senescence and apoptosis (8).

The enzyme O^6^-methylguanine-DNA methyltransferase (MGMT) removes guanine O^6^-methyl groups and restores DNA structure, and expression is thought to be related to temozolomide resistance. The expression of MGMT is in turn controlled by methylation of CpG islands on the MGMT promoter, suggesting that promoter methylation is a marker for temozolomide sensitivity. In addition, loss of mismatch repair (MMR) activity can confer resistance (7). The first aim of this study was to compare the proportion of temozolomide-sensitive cell lines in melanoma and glioblastoma. The second aim was to relate sensitivity in cell lines to MGMT expression and the third aim was to relate sensitivity to the methylation status of the MGMT promoter.

## Materials and Methods

### Cell lines and determination of IC50 values

Cell lines had been developed from surgical samples obtained with informed consent was obtained from patients using guidelines approved by the Northern Regional Ethics Committee and Auckland Area Health Board Ethics Committee. (1, 3). IC50 values (concentrations for 50% Inhibition of ^3^H-thymidine incorporation) were determined as previously described (3).

### Measurement of MGMT RNA synthesis

Standard curves were generated by serial dilutions of cDNA 1:10, 1:100, 1:1000 and 1:10,000. Cells were grown in T75 flasks unit ∼90% confluent. The media was removed, cells washed with 5 mL of PBS and immediately processed for RNA using the TRIzol LS reagent (Invitrogen) according to the manufacturer’s instructions. The RNA pellet was redissolved in 20 μL DEPC-treated water and quantified by spectrophotometry at 260 nm. The A260/280 ratio was used a measure of purity. To remove any contaminating DNA, the RNA samples were treated with DNAse using TURBO DNA-free (Ambion) protocol according to the manufacturer’s instructions. RNA (10 μg) was treated in a 50 μL reaction volume.

cDNA was produced using the Superscript III first strand synthesis system (Invitrogen). Two separate reactions were set up simultaneously, one with oligo(dT) and the other with random hexamers as the primers. Ten μL of DNA-free RNA (1 μg) was used as the template in each reaction. A -RT control was included for each sample. Following cDNA synthesis RNAse H was added to each reaction to remove any DNA/RNA duplexes. 2.5 μL of cDNA (diluted 1:100) was used for qPCR using the light cycler 480 with 0.5 μL of each primer (10 μmol/L), 5.0 μL of Absolute SYBR mix (2x) and water to a total volume of 10 μL. Each sample was analysed in triplicate and expression of MGMT compared to beta actin and GAPDH.

Light cycler protocol: Initial denaturation at 95 degrees C for 15 minutes, followed by 45 cycles of 95 degrees 10 sec, 61 degrees C 10 sec, 72 degrees C for 16 sec and 85 degrees C for 0 sec. The melt programme was as follows: 95 degrees C for 0 sec, 65 degrees C for 15 sec followed by cooling to 40 degrees for 30 sec. (This does not include ramp rates: if you need these I will need to find the archived files on the light cycler which might take some time).

Fluorescence acquisition was at 85 degrees C to exclude any primer dimers and then continuously measured during the pelt cycle for production of the melt curves. Data were analysed using the Relative Quantification software.

### Primers

MGMTfor CTTCACCATCCCGTTTTCCA

MGMTrev GATTGCCTCTCATTGCTCCT

Beta actin for GGGAAATCGTGCGTGACAT

Beta actin rev GAAGGAAGGCTGGAAGAGTG

GAPDHfor CGGGAAGCTTGTGATCCAATGG

GAPDHrev GGCAGTGATGGCATGGACTG

### Determination of MGMT methylation status

Bisulphite Conversion of DNA: The bisulphite conversion of DNA was performed using Zymo Research Kit – EZ DNA Methylation – Gold Kit (Catalog number D5005, Ngaio Diagnostics) according to the manufacturer’s instructions.

Analysis by Methylation-Specific PCR (MSP): DNA extraction was performed according to standard protocols described above. DNA methylation patterns in the CpG island of MGMT gene were determined by chemical modification of unmethylated cytosines to uracil, not the methylated cytosines to uracil. DNA extraction was performed according to standard protocols described above. DNA methylation patterns in the CpG island of *MGMT* gene were determined by chemical modification of unmethylated cytosines to uracil, not the methylated cytosines to uracil. Methylation-specific polymerase chain reaction (MSP) was performed with primers specific for either methylated or the modified unmethylated DNA, as previously described by (9). Primer sequences of *MGMT* for the unmethylated reaction were 5′-TTTGTGTTTTGATGTTTGTAGGTTTTTGT-3′ (MGMT_UF primer) and 5′-AACTCCACACTCTTCCAAAAACAAAACA-3′ (MGMT_UR primer) and for the methylated reaction were 5′-TTTCGACGTTCGTAGGTTTTCGC-3′ (MGMT_UF primer) and 5′-GCACTCTTCCGAAAACGAAACG-3′ (MGMT_UR primer) (9).

The PCR reaction for amplifying the bisulphite-converted DNA had the following components: 10 x PCR buffer (5 μl), primers (1.5 μl, forward and 1.5 μl reverse at 100 ng/ul each), 10mM dNTP (1 μl), 50mM MgCl_2_ (1.5 μl), 5 U/μl Platinum Taq polymerase (Life Technologies NZ Ltd - ThermoFisher Scientific, catalog 10966-034) (0.5 μl) and water (37 μl). The PCR conditions were 95 °C, 4 min; 35 cycles (95 °C, 30 s; 59 °C, 45 s; 72 °C, 45 s); 72 °C, 5 min; 4 °C hold. PCR products were run on a 4% low melting point agarose gel electrophoresis (with TBE buffer) for analysis.

## Results

### Sensitivity of cell lines to temozolomide

This was measured by inhibition of incorporation of radiolabelled thymidine into DNA. 40% of the melanoma lines and 24% of the glioblastoma lines showed sensitivity to temozolomide (Tables 1 and 2).

**Table 1.**
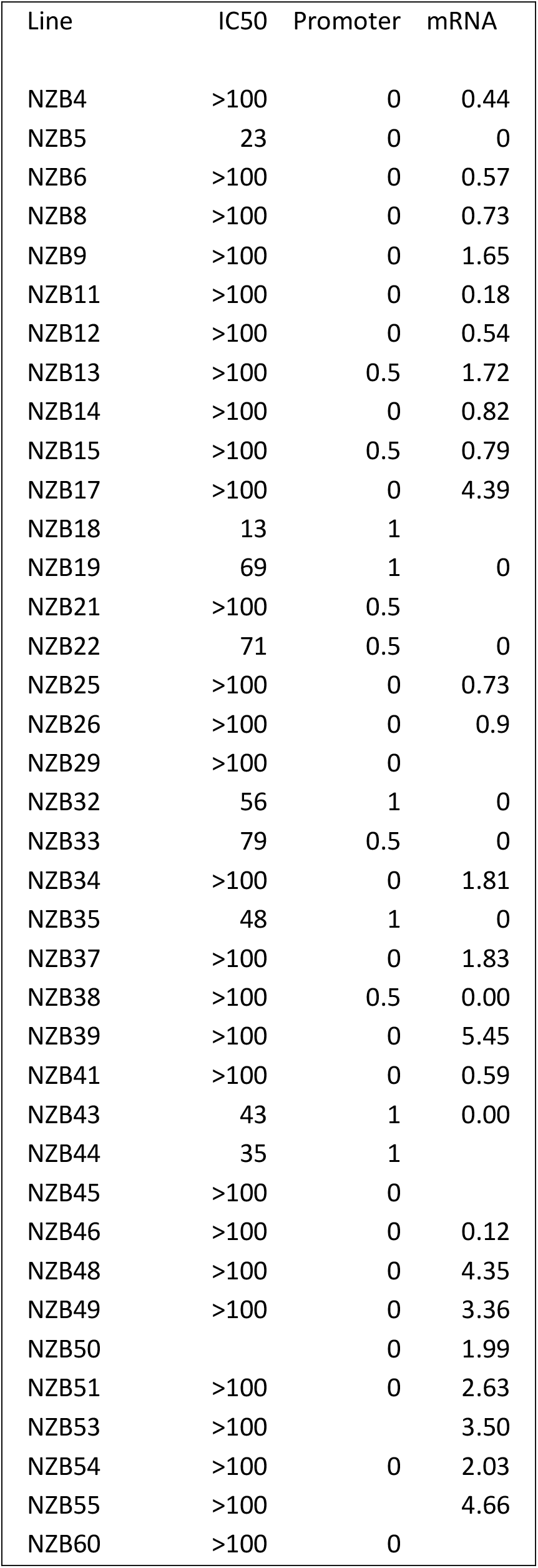
Brain tumour lines.

**Table 2.** Melanoma lines.

| Cell line | IC50 tem | Promoter | MGMT RNA |
| --- | --- | --- | --- |
| NZM1 | >100 | 0 | 0.10 |
| NZM2 | >100 | 0 | 1.00 |
| NZM3 | 33 | 1 | 0.00 |
| NZM4 | >100 | 0 | 1.43 |
| NZM5 | >100 | 0 | 0.65 |
| NZM6 | >100 | 0.5 | 0.38 |
| NZM7 | >100 | 0 | 2.49 |
| NZM9 | >100 |  | 0.00 |
| NZM10 | 10 | 0.5 | 0.00 |
| NZM11 | 19 | 1 | 0.00 |
| NZM12 | >100 | 0 | 1.32 |
| NZM13 | 22 | 0 | 0.02 |
| NZM14 | 75 | 1 | 0.00 |
| NZM15 | >100 | 0 |  |
| NZM16 | 81 |  | 0.00 |
| NZM18 | >100 | 0 | 0.23 |
| NZM19 | 53 | <0.5 | 0.01 |
| NZM20 | >100 | 0 | 0.16 |
| NZM21 | >100 | 0.5 | 0 |
| NZM22 | >100 | 0.5 |  |
| NZM23 | 150 | 0.5 | 0 |
| NZM24 | >100 | 0 | 0.06 |
| NZM25 | >100 |  | 0.05 |
| NZM27 | 120 |  | 0.00 |
| NZM28 | >100 | 0.5 | 0.58 |
| NZM29 | >100 | 0 | 0.28 |
| NZM30 | 77 | 1 | 0.00 |
| NZM31 | >100 | 0 |  |
| NZM32 | 12 |  | 0.09 |
| NZM33 | >100 | 0 | 0.12 |
| NZM34 | 35 | 1 | 0.00 |
| NZM35 | >100 | 0 |  |
| NZM36 | >100 | 0 | 0.04 |
| NZM40 | >100 | 0.5 | 0.01 |
| NZM41 | >100 | 0 | 0.02 |
| NZM43 | 80 | 0.5 | 0.20 |
| NZM44 | >100 | 0.5 |  |
| NZM45 | >100 | 0 | 1.15 |
| NZM46 | 60 | 0.5 | 0.00 |
| NZM47 | >100 | 0.5 | 0.01 |
| NZM48 | 71 | 0.5 | 0.01 |
| NZM49 | >100 | 0.5 |  |
| NZM50 | 11 | 0 | 0.03 |
| NZM52 | 85 | 0.5 |  |
| NZM53 | >100 | 0.5 | 0.34 |
| NZM54 | >100 | 0 | 1.08 |
| NZM55 | >100 | 0 | 1.69 |
| NZM56 | 92 | 0 | 0.00 |
| NZM58 | 59 | 0.5 | 0.01 |
| NZM59 | 37 | 1 | 0.00 |
| NZM60 | 26 |  | 0.00 |
| NZM61 | 89 | 0 | 0.00 |
| NZM62 | >100 | 0.5 | 0.03 |
| NZM63 | >100 | 0.5 | 1.28 |
| NZM65 | 82 | 0.5 |  |
| NZM66 | >100 | 0 |  |
| NZM67 | >100 | 0 |  |
| NZM68 | 11 | 1 |  |
| NZM69 | >100 | 0.5 | 0.00 |
| NZM71 | 76 | 0.5 |  |
| NZM72 | 27 | 0.5 |  |
| NZM79 | 67 | 0 |  |
| NZM83 | >100 | 0 |  |

### Expression of MGMT mRNA

This is shown in the Tables and in Fig. 1. Of the 79 cell lines analysed for MGMT mRNA expression, a good correlation was found to temozolomide resistance (p < 0.0001).

**Fig. 1.**
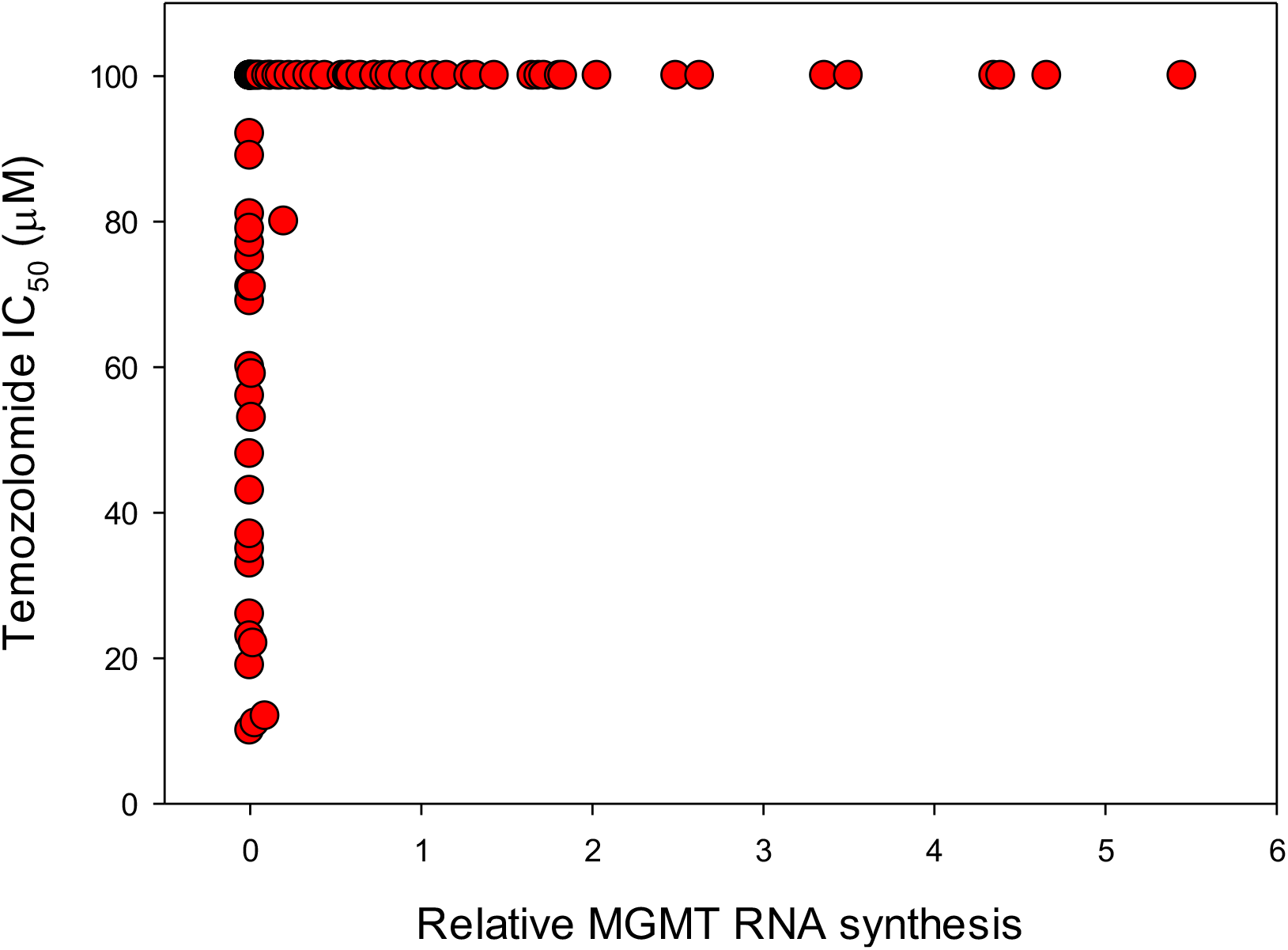
Relationship between MGMT expression and temozolomide sensitivity, as determined by inhibition of ^3^H-labelled thymidine incorporation (IC50 assays) among a series of NZM and NZB tumour lines. The highest drug concentration is depicted where 50% growth inhibition was not reached.

### MGMT promoter status of cell lines

This is shown in Tables 1 and 2. Of the 100 cell lines assessed, 12 showed methylation of both promoter sequences and were all temozolomide-sensitive; 50 lines showed no MGMT promoter methylation and 90% of these were temozolomide resistant; 30 lines showed partial promoter methylation and 53% of these were temozolomide resistant.

## Discussion

We show here that temozolomide inhibits the incorporation ^3^H-thymidine in a proportion of both melanoma and glioblastoma cell lines. In most but not all of these cases, inhibition is associated with methylation of the MGMT promoter, consistent with the hypothesis that of this promoter plays a major role in the epigenetic control of temozolomide sensitivity, with other mechanisms such as loss of mismatch repair playing a minor role (7). An important observation (Tables 1 and 2) is that while cell lines showing promoter methylation on both MGMT alleles are generally sensitive to temozolomide, some cell lines showed a mixed response, which might be explained by methylation affecting only one allele. MGMT promoter methylation status analysis by methylation-specific polymerase chain reaction, at least of the DNA sites measured in this study, is therefore not in itself sufficient to predict intrinsic sensitivity or resistance to temozolomide. This has been reported by others (10).

A feature of the results is the broad range of IC50 values obtained (Tables 1 and 2). Inhibition of ^3^H-thymidine incorporation, which measures proliferating cells in the S-phase of the cell division cycle (3, 11). In these studies the addition of unlabelled thymidine and fluorodeoxyuridine constrains cells to rely on exogenous thymidine rather than de novo synthesis (1).

In conclusion and within the limits of the assays, each of the cell lines appear to be homogeneous with respect to temozolomide sensitivity, MGMT mRNA expression and MGMT promoter methylation. This suggests that the tumours from which they were derived are also homogeneous, since there were no constraints imposed during their derivation. The conclusion is reinforced by the clinical observations that tumours were either sensitive or resistant to temozolomide. We would like to suggest that melanomas and gliomas undergo a type of clonal succession, in which a single stem cell populates the entire tumour for a given period, after which it is replaced by the progeny of a second stem cell, which may have a different sensitivity to temozolomide.

## Notes

**Funding** This work is supported by the Cancer Society New Zealand—Auckland and the Auckland Cancer Society Research Centre.

**Conflict of interests** The authors have no conflicts of interest to disclose.

### Competing Interest Statement

The authors have declared no competing interest.

